# Epstein–Barr virus exploits ER stress and TMX4-dependent nuclear envelope remodeling to enable capsid egress

**DOI:** 10.64898/2026.07.21.739849

**Authors:** Marika K. Kucińska, Tatiana Soldà, Diego Morone, Andrea Raimondi, Maurizio Molinari

**Affiliations:** Università della Svizzera italiana (USI), Faculty of Biomedical Sciences, Institute for Research in Biomedicine, CH-6500 Bellinzona, Switzerland; School of Life Sciences, École Polytechnique Fédérale de Lausanne, CH-1015 Lausanne, Switzerland

## Abstract

Herpesvirus capsids assemble within the nucleoplasm of infected cells in highly ordered icosahedral structures with a diameter of about 100nm. Despite the small distance between outer and inner nuclear membrane, which is fixed between the 25 and the 50nm by disulfide bonded LINC complexes, the capsid particles cross the barrier and are delivered into the cytoplasm, where viral particle assembly continues. How Epstein–Barr virus (EBV) overcomes the spatial constraints imposed by the narrow perinuclear space for nuclear egress of the viral capsids remains unclear. Here, we show that EBV exploits an ER-stress-responsive nuclear envelope (NE) remodeling pathway to promote capsid egress. Induction of EBV lytic replication activates the IRE1 branch of the unfolded protein response and triggers TMX4-dependent remodeling of the NE. Inhibition of IRE1 signaling or depletion of TMX4 prevents efficient redistribution of viral capsid proteins from the nucleus to the cytoplasm, causes accumulation of unused viral glycoprotein GP350 in Golgi-derived membranes, and markedly reduces production of infectious viral particles. Thus, EBV hijacks a host NE adaptation pathway to overcome a fundamental physical barrier during viral maturation.

## Introduction

EBV (human herpesvirus 4) infects about 95% of the adult population worldwide and establishes a lifelong latency ^1,2^. Viral re-activation that can occur upon immunosuppression or stress initiates the lytic replication cycle and the production of active viral particles ^3,4^. During the lytic phase, newly assembled nucleocapsids with a diameter of about 100nm are generated within the host cell nucleus. EBV capsids are too large to exit the nucleus through nuclear pore complexes via the canonical nucleoplasm-cytoplasm transport route. Instead, they first bud through the inner nuclear membrane (INM) into the perinuclear space and finally translocate across the outer nuclear membrane (ONM) to reach the cytoplasm. Subsequently, the viral particles acquire their envelope from Golgi-derived membranes and are eventually released from cells ^3,4^. Nuclear egress is mediated by the viral nuclear egress complex (NEC), comprising the EBV-encoded BFRF1 and BFLF2 proteins. The BFRF1– BFLF2 complex localizes to the INM, remodels the nuclear lamina and promotes membrane deformation to promote primary envelopment of nucleocapsids into the perinuclear space ^5–7^. This process is facilitated by the recruitment of host cell factors, including components of the ESCRT machinery ^8^ and cellular kinases that promote localized lamina disassembly. However, the precise contribution of host proteins is poorly defined. A major structural constraint is imposed by the linker of nucleoskeleton and cytoskeleton (LINC) complexes, which physically couple the INM and ONM through disulfide bond-dependent interactions between SUN proteins in the INM and NESPRIN proteins in the ONM ^9–12^. These complexes maintain the perinuclear space at only ∼25–50nm ^9,11,12^, substantially narrower than the diameter of EBV capsids, implying that successful nuclear egress requires local expansion of the nuclear envelope (NE). How this barrier is overcome has remained unknown. We recently identified the ER resident disulfide reductase TMX4 as a regulator of ER stress-induced NE remodeling ^13,14^. Upon mild ER stress induced by cyclopiazonic acid (CPA), a reversible inhibitor of the sarco/ER calcium-ATPase ^15^, TMX4 reduces the *inter*molecular disulfide bond linking SUN and NESPRIN proteins, thereby promoting LINC complexes disassembly and expanding the INM-ONM distance from 25-50nm to 100-200nm ^13,14^. Notably, EBV lytic replication activates the unfolded protein response (UPR), an ER stress pathway that promotes efficient viral assembly and production ^16–20^. These observations raised the possibility that EBV exploits the endogenous TMX4-dependent NE remodeling mechanism to overcome the spatial barrier imposed by LINC complexes during nuclear egress.

Here, we show that the EBV-induced ER stress and a TMX4-dependent pathway remodel the NE architecture and enables capsid egress from the nucleus. Our findings identify a host stress-adaptation pathway that is hijacked by EBV to facilitate herpesvirus maturation and reveal NE remodeling as a previously unrecognized host determinant of viral egress.

## Results

### EBV capsid egress requires extensive remodeling of the NE

To investigate how EBV capsids overcome the physical constraints imposed by the NE during their egress from the nucleoplasm into the cytoplasm, we used Human Embryonic Kidney 293 (HEK293) cells harboring the p2089 bacmid containing the complete genome of the EBV B95.8 strain engineered to express GFP (HEK293 EBV-eGFP cells) ^21–23^. This established cell line supports the full lytic cycle of EBV, including viral replication, capsid maturation, nuclear egress, and release of infectious virions. Transient expression of the immediate-early transcription factor BZLF1 triggers the lytic program, as shown by the induced expression of the viral capsid antigen p18, the major capsid protein (MCP), and the envelope glycoprotein GP350 transcripts (**Fig. 1a**). The lytic cycle is characterized by the induction of a mild UPR as shown by the enhanced detection of the spliced variant of the transcription factor XBP1 (sXBP1, **Fig. 1b**).

**Fig. 1:**
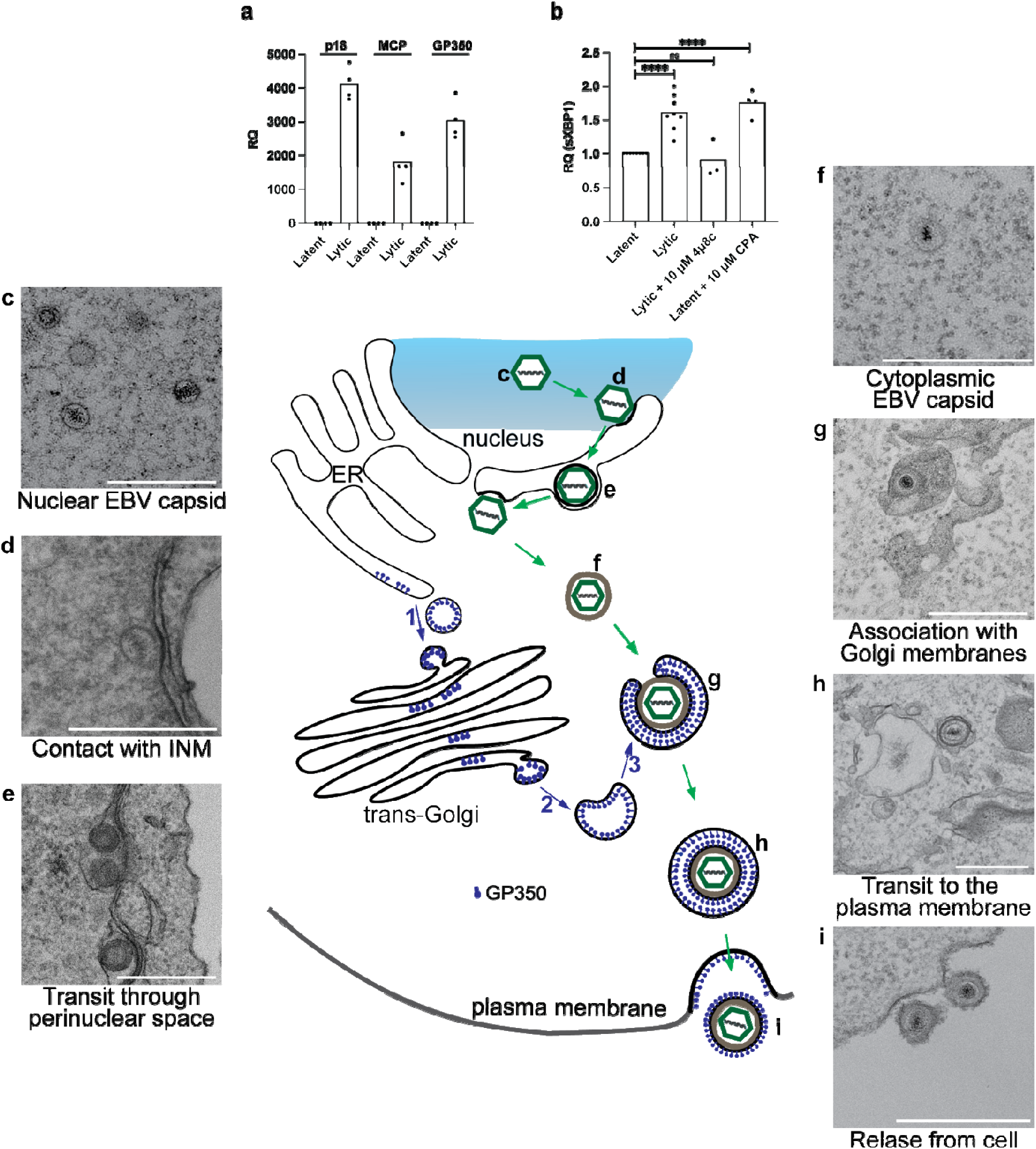
Analysis of EBV viral gene expression, ER stress responses and sequential stages of capsid maturation during EBV lytic cycle in HEK293 EBV-eGFP cells. **a)** Analysis of the viral gene transcripts p18, MCP, and GP350 48h after induction of the lytic phase as determined by qPCR. N=4 biological replicates. **b)** Analysis of sXBP1 transcripts during the latent phase (N=7), lytic phase (N=7), lytic phase in the presence of 10µM 4µ8c (N=3) or in the latent phase in the presence of 10µM CPA (N=4). An ordinary one-way ANOVA with Dunnett’s multiple comparisons test was performed. The mean bar is shown. ^ns^P>0.05, ****P<0.0001. **c-i)** Representative RT-TEM micrographs showing sequential stages of the EBV lytic cycle. Scale bars: 500nm.

To monitor the generation of virions upon lytic cycle induction, we performed room temperature-transmission electron microscopy (RT-TEM). Ultrastructural analyses captured successive stages of viral maturation, including nucleocapsid assembly within the nucleoplasm (**Fig. 1c**), docking of capsids at the INM (**Fig. 1d**), transit of capsids through the perinuclear space (**Fig. 1e**), release of capsids into the cytoplasm (**Fig. 1f**), cytoplasmic capsids associated with Golgi-derived membranes containing the viral glycoprotein GP350 (**Figs. 1g, 1h**), and release of mature virions from the plasma membrane (**Fig. 1i**). Release of the viral particles leaves a membrane layer containing GP350 at the plasma membrane of the infected cell (**Fig. 1**, schematics and panel **1i**). Notably, capsid transit across the NE was accompanied by pronounced local expansion and deformation of the perinuclear space (**Fig. 1e**). Given that the perinuclear space is normally maintained at a width of only ∼25–50nm, these observations indicate that EBV capsid egress is accompanied by extensive local remodeling of NE architecture to accommodate the passage of the nucleocapsids with an approximate diameter of 100nm.

### EBV lytic phase activates an ER stress pathway associated with NE remodeling

Activation of the UPR during EBV lytic phase supports production of viral proteins and virions ^16–20,24^. Because ER stress can induce TMX4-dependent remodeling of the NE ^13,14^, we investigated whether EBV activates this pathway to facilitate capsids’ egress from the nucleoplasm.

We monitored the activation of the IRE1 branch of the UPR in HEK293 EBV-eGFP cells following induction of the lytic cycle. Compared with latent cells, lytic replication increased splicing of the XBP1 transcripts, consistent with activation of a mild ER stress response (**Fig. 1b**, Latent *vs.* Lytic) ^16^. Splicing of XBP1 mRNA is operated by the ER stress transducer and RNase IRE1^25,26^. Consistently, the stress response induced during the EBV lytic cycle was abolished by the selective IRE1 RNase inhibitor 4µ8c ^27^, confirming pathway specificity (**Fig. 1b**, Lytic *vs.* 4µ8c). Induction of mild ER stress in latent cells using 10µM CPA, a condition previously shown to trigger TMX4-dependent NE remodeling ^14,28,29^, produced comparable levels of XBP1 splicing (**Fig. 1b**, Latent + 10µM CPA).

These findings demonstrate that the EBV lytic replication activates a mild IRE1-dependent ER stress response comparable in magnitude to that required for TMX4-mediated NE remodeling, supporting the hypothesis that EBV co-opts this host stress-adaptation pathway to promote nuclear egress.

### IRE1-dependent ER stress signaling enables EBV capsid egress from the nucleus

Mild ER stress triggers TMX4-dependent remodeling of the NE by promoting disassembly of LINC complexes and expansion of the perinuclear space ^13,14^. Because the EBV lytic replication induced an ER stress response of comparable magnitude (**Fig. 1b**) and was associated with local expansions of the NE to accommodate 100nm capsids during their transit through the perinuclear space (**Fig. 1e**), we tested whether activation of the IRE1 pathway is required for EBV capsid egress.

Lytic replication was induced in HEK293 EBV-eGFP cells in the presence or absence of the IRE1 RNase inhibitor 4μ8C ^27^, and the subcellular distribution of viral capsid antigen (VCA)-positive capsids was quantified by three-dimensional reconstruction of confocal image stacks. In control cells, VCA-positive capsids were detected in both the nucleus and the cytoplasm (**Fig. 2a**, VCA-positive capsids within the nucleoplasm are in red, VCA-positive capsids in the cytoplasm are in green, and quantification in **Figs. 2c, 2d**). In cells exposed to 10μM 4μ8C, VCA-positive capsids were detected within the nucleus, but not in the cytoplasm (**Figs. 2b-2d**), indicating that IRE1-dependent ER stress signaling is required for efficient EBV capsid egress from the nucleus.

**Fig. 2:**
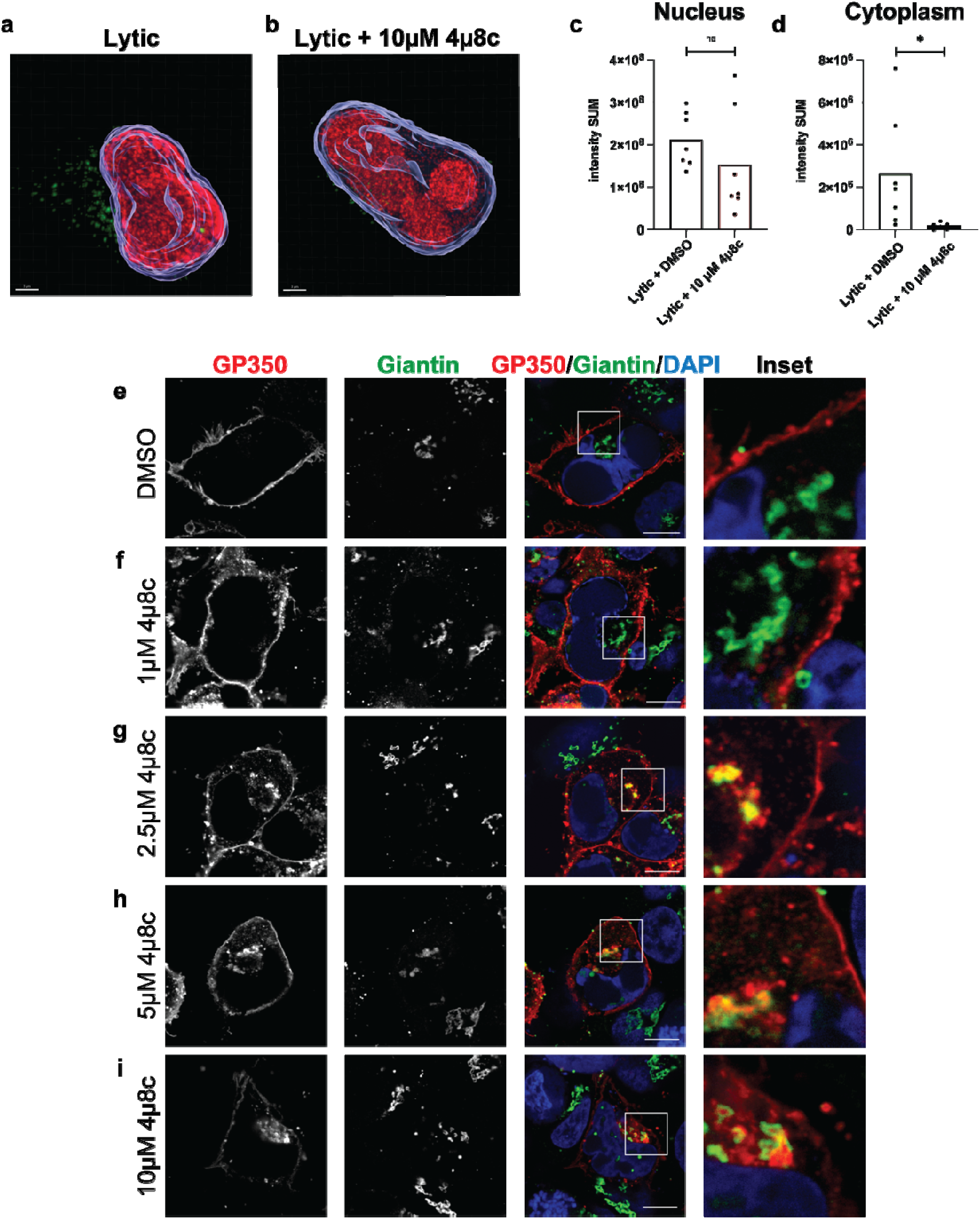
Effect of IRE1 inhibition on nuclear egress of EBV capsids. **a)** 3D-confocal image reconstruction of HEK293 EBV-eGFP VCA-positive cells 48h after lytic cycle induction in mock-treated cells **b)** Same as **(a)** in cells treated with 10µM 4µ8c during the lytic cycle. Scale bars: 3µm. **c)** Quantification of the intensity SUM of the VCA signal within the nucleus from (**a, b**). **d)** Same as (**c**) within the cytoplasm, n=7 cells per condition. The mean bar is shown. Unpaired two- tailed t-test, ^ns^P>0.05, *P<0.05. **e-i)** CLSM analysis of the EBV glycoprotein GP350 in cells mock-treated during 48h of lytic cycle or exposed to 4µ8c at 0, 1µM, 2.5µM, 5µM, 10µM. Scale bar: 10µm.

### IRE1 inhibition blocks virion maturation downstream of defective nuclear egress

We next examined whether impaired nuclear egress affects subsequent stages of EBV maturation by analyzing the localization of the EBV envelope glycoprotein GP350. GP350 is synthesized in the ER of infected cells (schematics in **Fig. 1**, arrow 1), traffics to the Golgi and Golgi derived vesicles (arrows 2, 3), where it meets and envelops the EBV capsids (schematics in **Fig. 1**, step **g**). In control cells, analyses by confocal laser scanning microscopy (CLSM) show that GP350 immunoreactivity is predominantly detected at the plasma membrane (**Fig. 2e**), consistent with efficient secondary envelopment, virion release, and retention of GP350-containing membrane at the cell surface following exocytosis (schematics and TEM micrographs, **Figs. 1g-1i**).

In contrast, inhibition of IRE1 signaling with 10µM 4μ8C, which caused nuclear retention of viral capsids (**Figs. 2b-2d**), resulted in a substantial redistribution of GP350 from the plasma membrane to the Golgi apparatus, where it extensively colocalized with the Golgi marker Giantin (**Fig. 2i**). These findings indicate that defective capsid export from the nucleus prevents efficient progression to the cytoplasmic maturation stages of the viral life cycle, leading to accumulation of GP350 within the Golgi compartment.

To determine whether this hampers the production of infectious virus, conditioned media from cells treated with increasing concentrations of 4μ8C during the lytic cycle (**Figs. 2e-2i**) were used to infect Raji cells (**Fig. 3** and Methods ^30–33^). Inhibition of IRE1 caused a dose-dependent reduction in the number of GFP-positive Raji cells (**Figs. 3a, 3b**). This demonstrated a corresponding decrease in the release of infectious EBV particles. Notably, this reduction closely correlated with the progressive loss of GP350 from the plasma membrane and its concomitant accumulation in the Golgi complex (**Figs. 2e-2i**). Together, these findings indicate that IRE1-dependent ER stress signaling is required for efficient EBV capsid nuclear egress and that disruption of this pathway blocks downstream virion maturation, ultimately reducing the production of infectious virus.

**Fig. 3:**
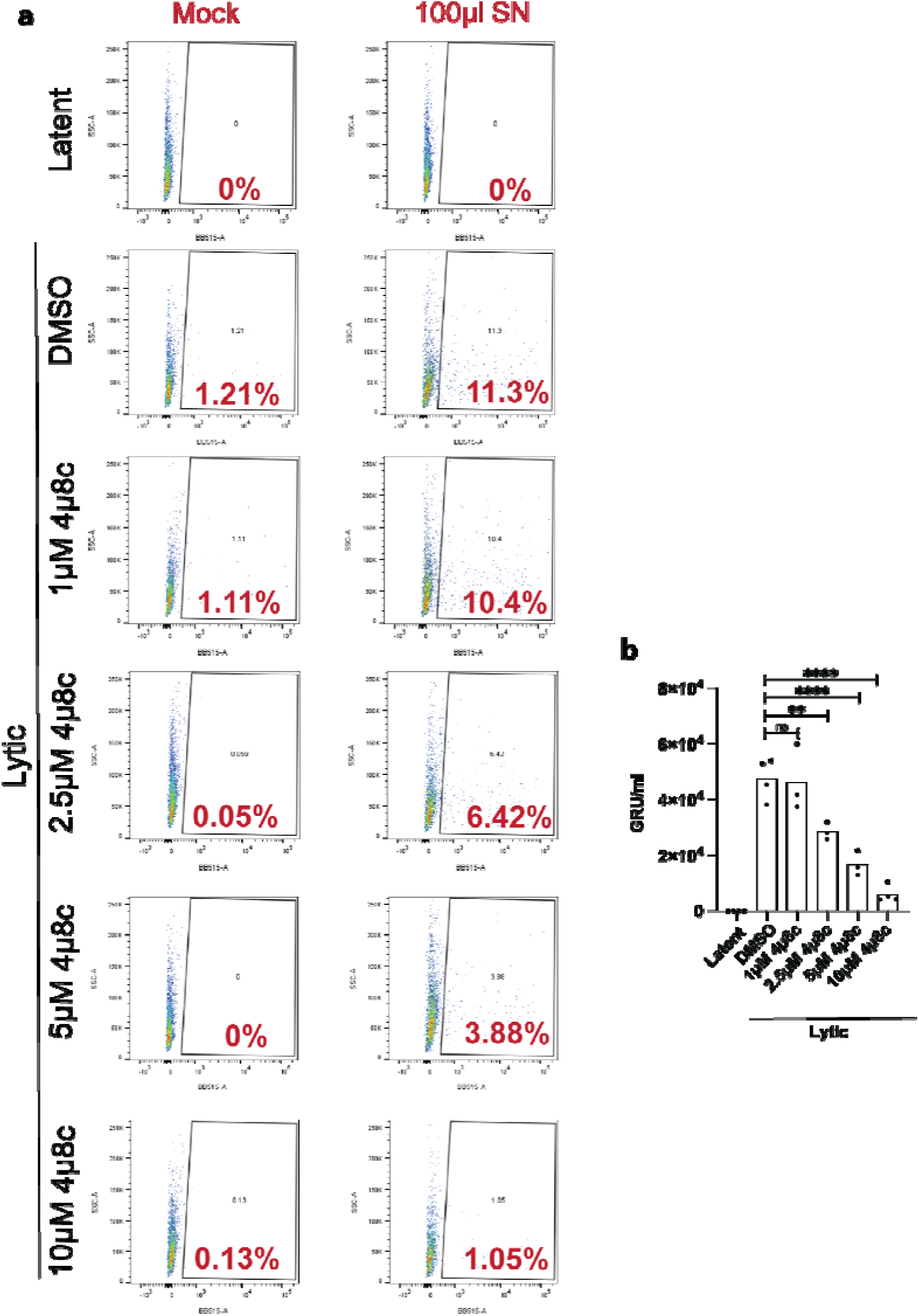
Assessment of infectious EBV particles produced by cells mock-treated or exposed to increasing concentrations of 4µ8c. **a)** Flow cytometry analysis of Raji cells mock incubated or incubated with 100µl of supernatant from HEK EBV-eGFP producer cells under the following conditions: latent phase, lytic induction, and lytic induction with 4µ8c at 1µM, 2.5µM, 5µM, 10µM. Raji cells were analyzed for GFP expression after 72h incubation with supernatants. **b)** Quantification of g een Raji units (GRU) per milliliter to determine the concentration of infectious EBV particles in panel **(a)**, N= at least 3 independent experiments. The mean bar is shown. An ordinary one-way ANOVA with Dunnett’s multiple comparisons test was performed, ^ns^P>0.05, ****P<0.0001.

### The ER-resident reductase TMX4 is required for EBV nuclear egress

To determine whether TMX4 mediates EBV nuclear egress, we depleted TMX4 by small interfering RNA (siTMX4, **Fig. 4a**) and compared the subcellular distribution of the VCA-positive capsids in lytically induced cells with that in cells transfected with a scrambled control siRNA (siSCR, **Fig. 4a**). Three-dimensional reconstruction of confocal image stacks showed that in siSCR control cells, VCA- positive capsids were distributed between the nucleus and the cytoplasm, consistent with efficient nuclear egress (red, **Figs. 4b**, quantification in **4d, 4e**). In contrast, TMX4 depletion caused a significant reduction in cytoplasmic capsids (**Figs. 4c, e**). Thus, loss of TMX4 phenocopied inhibition of IRE1 signaling, indicating that TMX4 is required for efficient export of EBV capsids from the nucleus. These findings are consistent with the established role of TMX4 in ER stress-induced NE remodeling ^13,14^ and support a model in which EBV exploits the host TMX4 pathway to expand the perinuclear space and facilitate capsid transit across the NE.

**Fig. 4:**
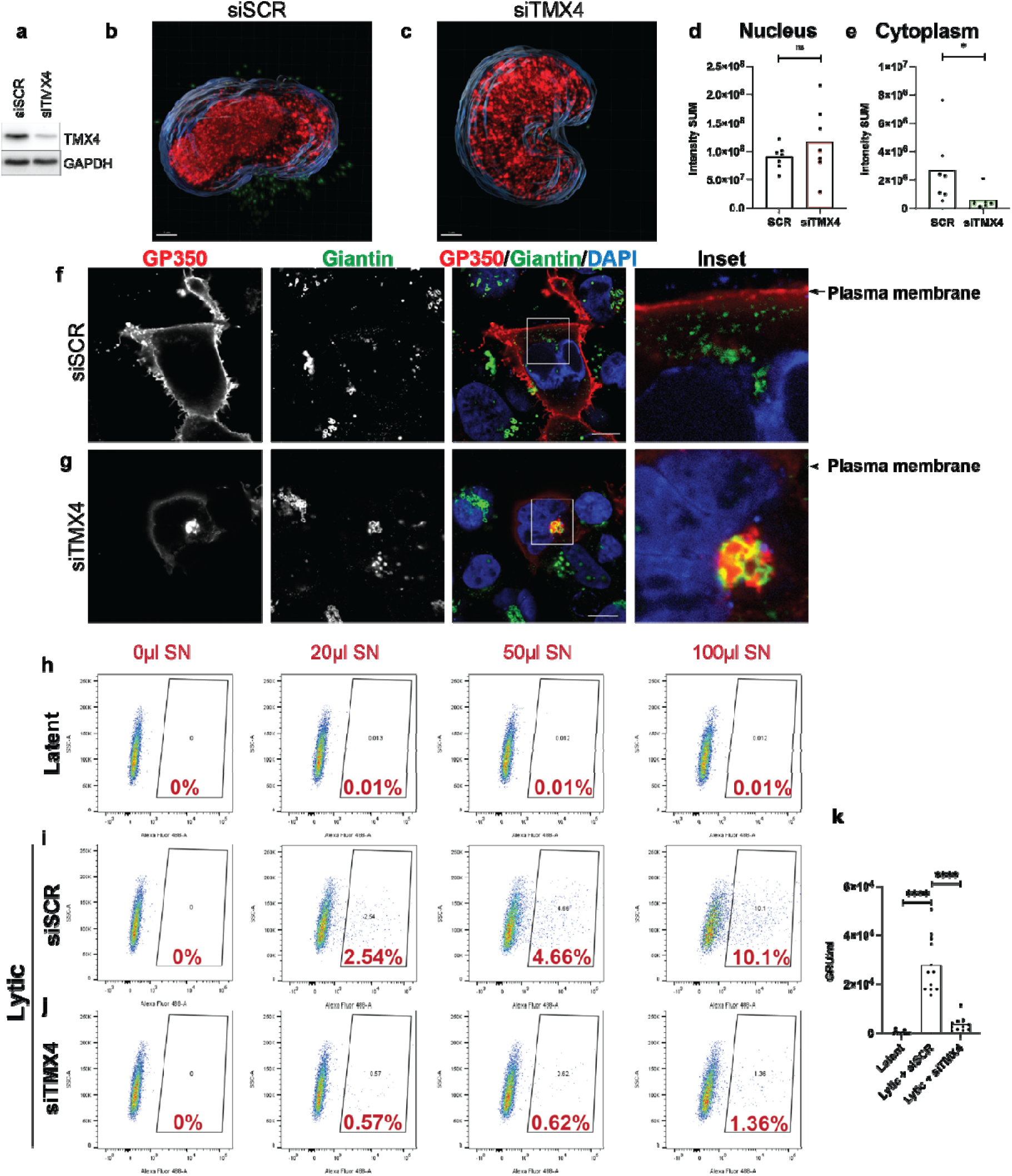
Effects of TMX4 silencing on EBV capsid protein and EBV viral glycoprotein localization, and infectious viral production. **a)** Western blot showing the level of endogenous TMX4 in cells expressing siSCR or siTMX4. **b)** 3D-confocal image reconstruction of VCA-positive capsids in HEK293 EBV-eGFP cells 48h after lytic cycle induction in mock-treated cells (siSCR). **c)** Same as **(b)** for siTMX4 cells. Scale bars: 3µm. **d)** Quantification of the intensity SUM of the VCA signal within the nucleus from (**b, c)** Same as (**d**) within the cytoplasm. n=7 cells per condition. The mean bar is shown. Unpaired, two-tailed t-test ^ns^P>0.05, *P<0.05. **f)** CLSM analysis of the EBV glycoprotein GP350 48h after induction of the lytic cycle in mock-treated (siSCR) cells. **g)** Same as (**f**) upon TMX4 silencing (siTMX4). Scale bars: 10µm. **h)** Flow cytometry analyses of Raji cells incubated with 0µl, 20µl, 50µl, and 100µl of the culture supernatant of cells in the latent phase. **i)** Same as **(h)** 48h after lytic cycle induction in mock-treated cells (siSCR). **(j)** Same as **(h)** in cells upon TMX4 silencing (siTMX4). **k)** Quantification of green Raji units (GRU) per milliliter to determine the concentration of infectious EBV particles. N= at least 3 independent experiments. The mean bar is shown. An ordinary one-way ANOVA with Dunnett’s multiple comparisons test was performed ****P<0.0001.

### TMX4-dependent nuclear egress coordinates virion maturation and infectious virus production

We next investigated whether impaired EBV nuclear egress following TMX4 silencing affected subsequent stages of viral maturation. In lytically induced control cells (siSCR), GP350 immunoreactivity was predominantly detected at the plasma membrane (**Fig. 4f**). In contrast, TMX4 silencing, which caused nuclear retention of viral capsids (**Figs. 4c-4e**), resulted in a marked redistribution of GP350 to the Golgi apparatus, where it extensively colocalized with the Golgi marker Giantin, with a corresponding reduction in plasma membrane staining (**Fig. 4g**). Again, loss of TMX4 phenocopied inhibition of IRE1 signaling that also resulted in the retention of unused GP350 in Giantin-positive Golgi structures (**Figs. 2f-2i**), thus confirming that impaired capsid export from the nucleus disrupts downstream virion maturation.

To determine whether these alterations compromised production of infectious virus, conditioned media from lytically induced siSCR- and siTMX4-treated cells were used to infect Raji cells, and infection efficiency was quantified by flow cytometric analysis of GFP-positive cells (**Figs. 4i, 4j, 4k**). Supernatants from TMX4-silenced cells exhibited a significantly reduced capacity to infect Raji cells (**Figs. 4j, 4k**) compared with controls (**Figs. 4i, 4k**), demonstrating that TMX4 silencing substantially hampers production of infectious EBV particles.

To validate the redistribution of GP350 at ultrastructural resolution, we performed immunoelectron microscopy. In lytically induced control cells, GP350 labeling was abundant at the plasma membrane (**Fig. 5a**), whereas only limited labeling was detected in Golgi-associated membranes (**Fig. 5b** and Inset). By contrast, TMX4 silencing substantially reduced GP350 labeling at the plasma membrane (**Fig. 5c**) and resulted in its prominent accumulation within the Golgi complex (**Fig. 5d** and Inset).

**Fig. 5:**
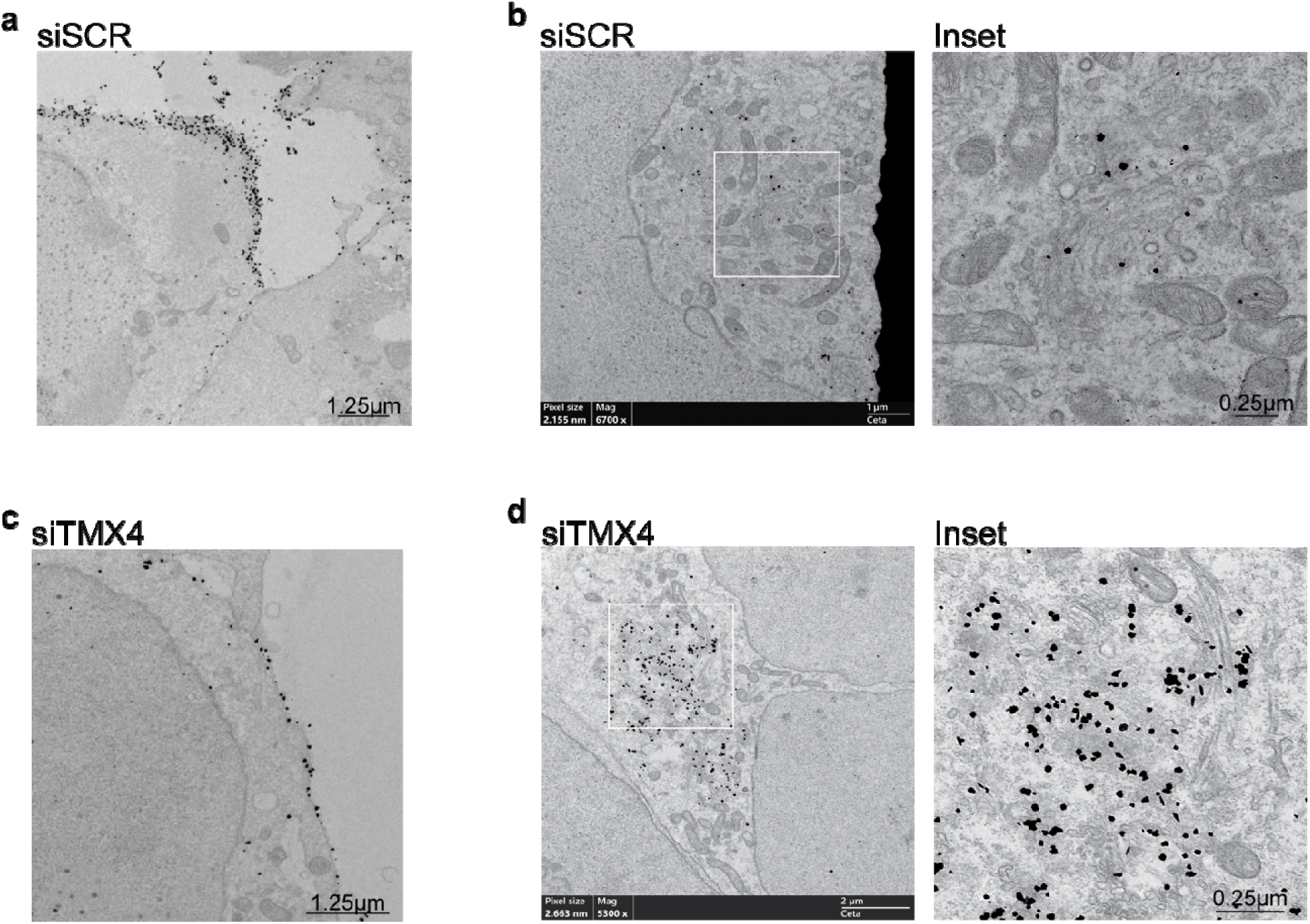
Subcellular distribution of EBV GP350. **a)** IEM micrographs showing immunogold-labeled EBV glycoprotein GP350, in HEK293 EBV-eGFP cells 48h post-lytic induction in the plasma membrane, mock-treated control (siSCR**) b**) Same as **(a)** in the Golgi apparatus. **(c)** Same as **(a)** upon TMX4 knockdown (siTMX4) **(d)** Same as **(b)** upon TMX4 knockdown.

These ultrastructural observations confirm that TMX4-dependent nuclear egress is required for efficient progression to the cytoplasmic stages of virion maturation.

## Discussion

Our study identifies a host stress-adaptation pathway that facilitates EBV nuclear egress by remodeling the NE. We show that the mild UPR elicited during the EBV lytic cycle ^16–20,24^ activates an IRE1- dependent signaling pathway that is required for efficient export of viral capsids from the nucleus. Mechanistically, our data support a model in which the ER-resident oxidoreductase TMX4 acts downstream of IRE1 to promote local expansion of the perinuclear space, thereby enabling the transit of nucleocapsids with a diameter of about 100nm across a compartment that is normally only 25-50nm wide ^9,11,12^. Pharmacological inhibition of IRE1 or depletion of TMX4 produced indistinguishable phenotypes, characterized by nuclear retention of capsids, failure of downstream virion maturation, intracellular accumulation of the envelope glycoprotein GP350 within Giantin-positive Golgi structures, and reduced production of infectious virus. Together, these findings reveal that EBV exploits a mild ER stress response to overcome a fundamental structural barrier imposed by the host cell.

Herpesvirus nuclear egress relies on the viral nuclear egress complex (NEC), composed in EBV of BFRF1 and BFLF2 ^5–7^, which remodels the inner nuclear membrane and nuclear lamina to promote primary envelopment of nucleocapsids. However, how the NE accommodates capsids that substantially exceed the physiological width of the perinuclear space has remained unresolved. Our ultrastructural analyses reveal pronounced local expansion of the NE surrounding transiting capsids, providing direct morphological evidence that large-scale remodeling of the host NE accompanies EBV nuclear egress. These observations suggest that, in addition to NEC-mediated membrane deformation, successful nuclear escape requires host mechanisms that transiently increase the spacing between the inner and outer nuclear membranes.

Our findings identify the IRE1-TMX4 axis as a candidate mechanism for this remodeling. We recently demonstrated that mild ER stress activates TMX4 to reduce disulfide bonds linking SUN and NESPRIN proteins within LINC complexes, thereby uncoupling the inner and outer nuclear membranes and locally expanding the perinuclear space ^13,14^. The observation that EBV induces an ER stress response of similar magnitude, together with the phenocopy obtained by IRE1 inhibition and TMX4 depletion, strongly supports the idea that the virus co-opts this endogenous pathway to facilitate nuclear egress.

More broadly, our work uncovers a previously unrecognized role for EBV-induced UPR signaling in regulating NE architecture during viral infection. Rather than representing a passive consequence of the high secretory burden associated with lytic replication, mild ER stress emerges as an active host process exploited by EBV to coordinate nuclear egress with downstream virion maturation. Given that nuclear egress is a conserved feature of all herpesviruses, it will be important to determine whether remodeling of the NE through the IRE1-TMX4 pathway represents a broader strategy used by this virus family. Such a mechanism would identify host pathways controlling nuclear membrane organization as attractive antiviral targets that may be less susceptible to the emergence of viral resistance than therapies directed against viral proteins.

## Methods

### Cell lines

HEK293 cells containing the recombinant EBV strain B95-8 DNA in the form of a bacmid, p2089, maintained under Hygromycin selection and expressing the eGFP protein, were kindly provided by Dr. C. Münz Laboratory, University Zurich and are described in ^33^. They were grown in RPMI medium with 10% FBS, hygromycin and Pen/Strep. Raji cells were kindly provided by prof. Münz. They were grown in RPMI with 10% FBS, and Pen/Strep.

### EBV lytic cycle induction

HEK293 EBV-eGFP cells were seeded in 12-well or 3.5cm^2^ dishes and, the day after, were transfected with 0.4μg or 1μg of BZLF1 (p509), respectively, using PEI MAX transfection reagent (6μl of PEI Max 1mg/ml per 1μg of DNA) in Opti-MEM and incubated for 20min RT.

### RNA silencing

RNA interference was performed in the HEK EBV-eGFP 12h after lytic cycle induction.

Cells were transfected using JetPrime transfection reagent, (PolyPlus) following the manufacturer’s protocol with scrambled small interfering RNA or small interfering RNA against TMX4 (sense strand: 5’-GAUUCAGAAUGGGAGGCUUtt-3’antisense strand 5’-AAGCCUCCCAUUCUGAAUCag-3’ 100nM per well in 12MW, Ambion, Life Technology). Cells were analyzed by immunofluorescence and biochemical analyses after 36h of silencing.

### Cell lysis and Western blot

Following the respective treatments, cells were washed with ice-cold PBS containing 20 mM N-ethylmaleimide (NEM). Then, cells were lysed in RIPA buffer (1% Triton X-100, 0.1% SDS, 0.5% sodium deoxycholate in HBS, pH 7.4) containing 20 mM NEM and protease inhibitor cocktail and collected by scraping, incubated on ice for 20 min. Post-nuclear supernatant (PNS) was collected by centrifugation at 10,000 × g, 4°C for 10 min. The PNS was denatured and reduced in 10 mM dithiothreitol (DTT)-containing sample buffer and heated at 95°C for 5min and subjected to polyacrylamide gel electrophoresis (SDS-PAGE). Proteins were transferred to the PVFD membrane using the Trans-Blot Turbo Transfer System (Bio-Rad). Afterwards, the membrane was blocked with 8% (w/v) non-fat dry milk for 10min in Tris-buffered saline containing 1% Tween 20 (TBS-T) and stained with the primary antibodies for 90min diluted in TBS-T under agitation at RT. Then, HRP-conjugated secondary antibodies or Protein A HRP-conjugated were applied for 45min, diluted in TBS- T at RT. The protein bands were visualized using the WesternBright Quantum (Witec) and signals were captured with the FusionFX7 VILBER chemiluminescence imaging system (Witec) using Fusion FX7 Edge software.

### Virus infectivity determination by flow cytometry

Supernatants containing EBV that encodes an eGFP reporter gene, produced in HEK293 EBV-eGFP cells, were harvested, centrifuged at 2’000g for 10min and filtered through 1.2μm filter. Afterwards, supernatants were titrated and incubated with 40 000 Raji cells in 96 MW for 72h hours, resulting in intracellular eGFP expression in successfully infected Raji cells. Cells were washed twice in PBS-/- (without Ca++ and Mg++) and incubated with live/dead staining for 15min, following one PBS-/- wash, resuspended in MACS buffer (PBS -/- with 2% FBS and 2mM EDTA) and run on FACSymphony^TM^ A5/A3 or BD LSRFortessa. The infection level was determined by measuring the eGFP-positive cell’s percentage, with subsequent data analysis performed in FlowJo. Live/dead staining was performed with LIVE/DEAD Fixable Dead Cell Stain Kit (ThermoFisher) or with DAPI (Sigma) to exclude dead cells following the manufacturer’s protocol. Quantification of viral infectivity was reported as Green Raji Units (GRU) per milliliter. Formula: number of seeded Raji cells x (% of GFP-positive cells/100) / virus volume (mL) = virus particles/ml.

### Confocal laser scanning microscopy (CLSM)

HEK293 EBV-eGFP cells were seeded on alcian blue-treated glass coverslips and treated as indicated in the figure’s legend. Cells were fixed in 3.7% formaldehyde diluted in PBS for 20 min in RT. For GP350, cells were permeabilized with 0.05% saponin, 10% goat serum, 10mM HEPES and 15mM glycine for 15min. Cells were incubated with primary antibodies for 2h, washed twice in PBS, and incubated with secondary antibody for 45min in RT. After incubation time, cells were washed three times in PBS and once in water. Subsequently, coverslips with cells were mounted with Vectashield with DAPI (Vector Laboratories). For VCA, cells were permeabilized in 0.2% Triton in PBS for 20min in RT. After washing once in PBS, blocking was performed in 1% FBS for 1h in RT, followed by one wash in PBS. Cells were incubated with primary antibodies for 2h in RT, washed twice in PBS. Subsequently, cells were incubated with secondary antibodies for 45min in RT. Cells were washed three times in PBS and once in water. Afterwards, cells were mounted with Vectashield with DAPI (Vector Laboratories). Leica TCS SP5 microscope with a Leica HCX PL APO lambda blue 63.0 x 1.40 OIL UV objective was used to acquire microscope images. Confocal z-stacks were collected using a pinhole set to 0.8 U. Upon collection of all optical sections from top to bottom of the cell, the analysis of the acquired stacks was performed with Imaris 10.2.0.

### Quantification of VCA SUM intensity inside and outside the nucleus in Imaris 10.2.0

Imaris 10.2.0 software was used to reconstruct the 3D model from confocal image Z-stacks. The face surface tool was used to manually segment the nucleus and to automatically define the second surface, the cytoplasm. The statistical analysis of the intensity sum of VCA inside and outside the nucleus was performed in the Imaris software.

### RT-TEM

HEK EBV-eGFP were plated on alcian blue-coated glass coverslips and treated as described in the manuscript and initially fixed by adding fixative solution (4% PFA EM grade, 5% GA in Na-cacodylate buffer 0.1M, pH 7.4) directly to the culture media for 20 min in RT. The fixative was then removed, and cells were further incubated with a single-strength fixative solution (2% PFA, 2.5% GA in Na-cacodylate buffer 0.1M, pH 7.4) for 3h in RT. Following multiple washes in Na-cacodylate buffer, cells were post-fixed with 1% osmium tetroxide and 1.5% potassium ferricyanide in 0.1M Na-cacodylate buffer on ice. Samples were then rinsed with distilled water and subjected to enbloc staining with 0.5% uranyl acetate in distilled water overnight at 4 °C in the dark. After additional washes in distilled water, specimens were dehydrated through a graded ethanol series, embedded in Epon resin, and polymerized at 60 °C for 48h. Ultrathin sections (70-90nm) were prepared using an ultramicrotome (UC7, Leica Microsystems, Vienna, Austria), counterstained with uracyl acetate and Sato’s lead solution, and examined using a Talos L120C transmission electron microscope (FEI, Thermo Fisher Scientific) operated at 120 kV. Images were captured with a Ceta CCD camera (FEI, Thermo Fisher Scientific).

### IEM

HEK EBV-eGFP cells were fixed in Periodate-lysine-paraformaldehyde solution for 2h at RT. The cells were washed 3 times in PBS, and afterwards incubated with 50mM glycine for 10 min and permeabilized in saponin solution (0.1% saponin, 0.2% BSA, 5% goat serum, 50mM NH4Cl, 20mM PO_4_ buffer, 150mM NaCl) for 1h at RT. The primary antibody (GP350, 1:25) and nanogold-labeled secondary antibody (Fab anti-mouse 1:100, Nanoprobes) staining were executed in saponin solution at RT. Subsequently, cells were re-fixed in 1% GA for 30 min, and gold enhancement solution (Nanoprobes) was used for nanogold enlargement following the manufacturer’s protocol. Cells were then post-fixed with osmium tetroxide, embedded in Epon, and sectioned in ultrathin slices. Then, the sample contrasting was performed with uranyl acetate and lead citrate and acquired with Talos L120C TEM (FEI, Thermo Fisher Scientific) operating at 120 kV. Images were captured with a Ceta CCD camera (FEI, Thermo Fisher Scientific).

### Protocol to induce ER stress

HEK EBV-eGFP were exposed for 48h to 10µM CPA to induce ER stress ^14^.

### Protocol to inhibit IRE1α RNase activity

HEK EBV-eGFP cells were exposed for 48h to 1µM, 2.5µM, 5µM, 10µM and 25µM of 4µ8c (Merck) to inhibit IRE1α RNase activity during EBV lytic cycle.

### qRT-PCR

HEK EBV-eGFP cells were subjected to RNA extraction with GenElute Mammalian Total RNA Miniprep Kit (Sigma) following the manufacturer’s protocol. For the cDNA synthesis 1µg of RNA, dNTPs, oligo(dT)and the SuperScript II reverse transcriptase (ThermoFisher) were used, following the manufacturer’s protocol. For qRT-PCR, 1µl of cDNA was mixed with 1µl of 10µM forward and reverse primer mix, 10µl of Power SYBR Green PCR Master Mix (Selleckchem), 0.4µl of reference ROX dye and 7.6µl of sterile milliQ water. Samples were loaded as triplicates, vortexed and centrifuged in a 96-well reaction plate. Quantitative real-time PCR was performed using the QuantStudio^TM^3 Real-Time PCR System. Analysis of the data were performed using the QuantStudio^TM^ Design & Analysis Software v.1.5.5.

Beta Actin (fwd: 5’-CTTCCTGGGCATGGAGTCCT-3’; rev: 5’-GGAGCAATGATCTTGATCTT-3’); sXBP1 (fwd: 5’-CTGAGTCCGCAGCAGGTG-3’; rev: 5’-TGGCTGGATGAAAGCAGATT-3’); VCA-p18 (BFRF3) (fwd: 5’-TGAACCAGAATAATCTCCCCAATG-3’; rev: 5’-GCCGAGGCACCC CAAAAGTC-3’); MCP (BcLF1) (fwd: 5’-GGATGCCGCCTATGAATACC-3’; rev: 5’- CTGTACTCG TTGACCATGTTGT-3’); GP350 (BLLF1) (fwd: 5’- GCAACAAGTAAGCCTGGAATCT-3’; rev: 5’-CCTCACTACTGCCGTTATATTGG 3’).

### Statistical analysis

Graphs and statistical analyses were performed using GraphPad Prism 11. In this paper, one-way ANOVA with Dunnett’s multiple comparisons test and unpaired, two-tailed t-test were used to assess statistical significance. A P-value>0.05 was considered as statistically no significant (ns). Statistical significance was defined as *P<0.05, **P<0.01, ***P<0.001, and ****P<0.0001.

## Acknowledgments

We thank the members of Molinari’s lab for discussions, and critical reading of the manuscript. We thank Christian Münz and Maria Pena Francesch for reagents and discussion. We would like to thank David Jarossay for FACS technical support.

## Data availability

The source data are provided with this paper as a Source Data file.

## Author contribution

M.K.K and T.S. designed and performed the experiments, and the biochemical, imaging, transcriptional and quantitative analyses, analyzed data, and reviewed the manuscript; D.M. assisted in CSLM analyses. A.R. performed the analyses with the RT-TEM. M.M. designed the study, analyzed data, and wrote the manuscript. All the authors discussed the results and the manuscript.

The authors declare no competing interests.

## References

1 Wong, Y., Meehan, M. T., Burrows, S. R., Doolan, D. L. & Miles, J. J. Estimating the global burden of Epstein-Barr virus-related cancers. J Cancer Res Clin Oncol 148, 31–46 (2022). 10.1007/s00432-021-03824-y

2 Dunmire, S. K., Verghese, P. S. & Balfour, H. H., Jr. Primary Epstein-Barr virus infection. J Clin Virol 102, 84–92 (2018). 10.1016/j.jcv.2018.03.001

3 Dohner, K., Serrero, M. C., Viejo-Borbolla, A. & Sodeik, B. A Hitchhiker’s Guide Through the Cell: The World According to the Capsids of Alphaherpesviruses. Annu Rev Virol 11, 215–238 (2024). 10.1146/annurev-virology-100422-022751

4 Nanbo, A. Current Insights into the Maturation of Epstein-Barr Virus Particles. Microorganisms 12, 1–11 (2024). 10.3390/microorganisms12040806

5 Thorsen, M. K., Draganova, E. B. & Heldwein, E. E. The nuclear egress complex of Epstein- Barr virus buds membranes through an oligomerization-driven mechanism. Plos Pathog 18, e1010623 (2022). 10.1371/journal.ppat.1010623

6 Dai, Y. C. et al. The Novel Nuclear Targeting and BFRF1-Interacting Domains of BFLF2 Are Essential for Efficient Epstein-Barr Virus Virion Release. J Virol 94, e01498–01419 (2020). 10.1128/JVI.01498-19

7 Lee, C. P. & Chen, M. R. Conquering the Nuclear Envelope Barriers by EBV Lytic Replication. Viruses 13, 702 (2021). 10.3390/v13040702

8 Lee, C. P. et al. The ESCRT machinery is recruited by the viral BFRF1 protein to the nucleus- associated membrane for the maturation of Epstein-Barr Virus. Plos Pathog 8, e1002904 (2012). 10.1371/journal.ppat.1002904

9 Sosa, B. A., Rothballer, A., Kutay, U. & Schwartz, T. U. LINC Complexes Form by Binding of Three KASH Peptides to Domain Interfaces of Trimeric SUN Proteins. Cell 149, 1035–1047 (2012). 10.1016/j.cell.2012.03.046

10 Padmakumar, V. C. et al. The inner nuclear membrane protein Sun1 mediates the anchorage of Nesprin-2 to the nuclear envelope. Journal of Cell Science 118, 3419–3430 (2005). 10.1242/jcs.02471

11 Crisp, M. et al. Coupling of the nucleus and cytoplasm: role of the LINC complex. Journal of Cell Biology 172, 41–53 (2006). 10.1083/jcb.200509124

12 Sosa, B. A., Kutay, U. & Schwartz, T. U. Structural insights into LINC complexes. Current Opinion in Structural Biology 23, 285–291 (2013). 10.1016/j.sbi.2013.03.005

13 Kucinska, M. K. & Molinari, M. Control of nuclear envelope dynamics during acute ER stress by LINC complexes disassembly and selective, asymmetric autophagy of the outer nuclear membrane. Autophagy 20, 1–3 (2024). 10.1080/15548627.2023.2299123

14 Kucinska, M. K. et al. TMX4-driven LINC complex disassembly and asymmetric autophagy of the nuclear envelope upon acute ER stress. Nat Commun 14, 1–20 (2023). 10.1038/s41467-023-39172-3

15 Pirot, P. et al. Global profiling of genes modified by endoplasmic reticulum stress in pancreatic beta cells reveals the early degradation of insulin mRNAs. Diabetologia 50, 1006–1014 (2007). 10.1007/s00125-007-0609-0

16 Johnston, B. P. & McCormick, C. Herpesviruses and the Unfolded Protein Response. Viruses 12, 1–26 (2019). 10.3390/v12010017

17 Bhende, P. M., Dickerson, S. J., Sun, X., Feng, W. H. & Kenney, S. C. X-box-binding protein 1 activates lytic Epstein-Barr virus gene expression in combination with protein kinase D. J Virol 81, 7363–7370 (2007). 10.1128/JVI.00154-07

18 Chen, L. W. et al. The Epstein-Barr Virus Lytic Protein BMLF1 Induces Upregulation of GRP78 Expression through ATF6 Activation. Int J Mol Sci 22, 1–20 (2021). 10.3390/ijms22084024

19 Sun, C. C. & Thorley-Lawson, D. A. Plasma cell-specific transcription factor XBP-1s binds to and transactivates the Epstein-Barr virus BZLF1 promoter. J Virol 81, 13566–13577 (2007). 10.1128/JVI.01055-07

20 Taylor, G. M., Raghuwanshi, S. K., Rowe, D. T., Wadowsky, R. M. & Rosendorff, A. Endoplasmic reticulum stress causes EBV lytic replication. Blood 118, 5528–5539 (2011). 10.1182/blood-2011-04-347112

21 Delecluse, H. J., Hilsendegen, T., Pich, D., Zeidler, R. & Hammerschmidt, W. Propagation and recovery of intact, infectious Epstein-Barr virus from prokaryotic to human cells. Proc Natl Acad Sci U S A 95, 8245–8250 (1998). 10.1073/pnas.95.14.8245

22 Pena-Francesch, M. et al. The autophagy machinery interacts with EBV capsids during viral envelope release. Proc Natl Acad Sci U S A 120, e2211281120 (2023). 10.1073/pnas.2211281120

23 Feederle, R., Bartlett, E. J. & Delecluse, H. J. Epstein-Barr virus genetics: talking about the BAC generation. Herpesviridae 1, 1–13 (2010). 10.1186/2042-4280-1-6

24 Sun, C. C. & Thorley-Lawson, D. A. Plasma cell-specific transcription factor XBP-1s binds to and transactivates the Epstein-Barr virus BZLF1 promoter. J Virol 81, 13566–13577 (2007). 10.1128/Jvi.01055-07

25 Yoshida, H., Matsui, T., Yamamoto, A., Okada, T. & Mori, K. XBP1 mRNA is induced by ATF6 and spliced by IRE1 in response to ER stress to produce a highly active transcription factor. Cell 107, 881–891 (2001).

26 Calfon, M. et al. IRE1 couples endoplasmic reticulum load to secretory capacity by processing the XBP-1 mRNA. Nature 415, 92–96 (2002).

27 Cross, B. C. et al. The molecular basis for selective inhibition of unconventional mRNA splicing by an IRE1-binding small molecule. Proc Natl Acad Sci U S A 109, 869–878 (2012). 10.1073/pnas.1115623109

28 Fumagalli, F. et al. Translocon component Sec62 acts in endoplasmic reticulum turnover during stress recovery. Nat Cell Biol 18, 1173–1184 (2016). 10.1038/ncb3423

29 Loi, M., Raimondi, A., Morone, D. & Molinari, M. ESCRT-III-driven piecemeal micro-ER-phagy remodels the ER during recovery from ER stress. Nat Commun 10, 5058 (2019). 10.1038/s41467-019-12991-z

30 Pulvertaft, J. V. Cytology of Burkitt’s Tumour (African Lymphoma). Lancet 1, 238–240 (1964). 10.1016/s0140-6736(64)92345-1

31 Chen, Y. F. et al. Assessing the fitness of Epstein-Barr virus following its reactivation. J Virol 99, e0062625 (2025). 10.1128/jvi.00626-25

32 Polack, A., Delius, H., Zimber, U. & Bornkamm, G. W. Two deletions in the Epstein-Barr virus genome of the Burkitt lymphoma nonproducer line Raji. Virology 133, 146–157 (1984). 10.1016/0042-6822(84)90433-1

33 Delecluse, H. J., Hilsendegen, T., Pich, D., Zeidler, R. & Hammerschmidt, W. Propagation and recovery of intact, infectious Epstein-Barr virus from prokaryotic to human cells. Proc Natl Acad Sci U S A 95, 8245–8250 (1998). 10.1073/pnas.95.14.8245

